# A large reverse-genetic screen identifies numerous regulators of testis nascent myotube collective cell migration and collective organ sculpting

**DOI:** 10.1101/2024.10.10.617659

**Authors:** Maik C. Bischoff, Jenevieve E. Norton, Erika A. Munguia, Noah J. Gurley, Sarah E. Clark, Rebecca Korankye, Emmanuel Addai Gyabaah, Taino Encarnacion, Christopher J. Serody, Corbin D. Jones, Mark Peifer

## Abstract

Collective cell migration is critical for morphogenesis, homeostasis, and wound healing. During development migrating mesenchymal cells form tissues that shape some of the body’s organs. We have developed a powerful model for examining this, exploring how Drosophila testis nascent myotubes migrate onto the testis during pupal development, forming the muscles that ensheath it and also creating its characteristic spiral shape. To define genes that regulate this process, we have carried out RNAseq to define the genes expressed in myotubes during migration. Using this dataset, we curated a list of 131 ligands, receptors and cytoskeletal regulators, including all Rho-family GTPase GAPs and GEFs, as candidates. We then used the GAL4/UAS system to express 279 shRNAs targeting these genes, using the muscle specific driver dMef2>GAL4, and examined the adult testis. We identified 29 genes with diverse roles in testis morphogenesis. Some have phenotypes consistent with defects in collective cell migration, while others alter testis shape in different ways, revealing some of the underlying logic of testis morphogenesis. We followed up one of these genes in more detail—that encoding the Rho-family GEF dPix. dPix knockdown leads to a drastic reduction in migration and a substantial loss of muscle coverage. Our data suggest different isoforms of dPix play distinct roles in this process, reveal a role for its protein partner Git. We also explore whether cdc42 activity regulation or cell adhesion are among the dPix mechanisms of action. Together, our RNAseq dataset and genetic analysis will provide an important resource for the community to explore cell migration and organ morphogenesis.

## Introduction

Cell migration is critical for both normal development and organogenesis, and for tissue homeostasis. Some cells, like leukocytes, migrate individually, but many cells migrate collectively (Scarpa and Mayor, 2016; Mishra *et al*., 2019). These include cells that migrate as epithelial sheets, like *Drosophila* follicle cells during oogenesis or cells closing wounds in mammals, while other cells remain more loosely linked by cadherin-based junctions but take on more mesenchymal characteristics like vertebrate neural crest cells or some collectively migrating cancer cells (Mishra *et al*., 2019). These diverse modes of migration reflect differences in the underlying cell machinery, both that mediating the cytoskeletal events that mediate motility and in the guidance cues that attract or repel cells. One key challenge for our field is to define these underlying mechanisms.

Much of what we know about the cell migration machinery comes from studying cells in vitro, either in simplified two-dimensional settings, often with fibroblasts as a model, or in more complex three-dimensional models. These systems offer advantages, including relative ease of manipulation and access to high resolution imaging. However, we ultimately need to understand how cells migrate in their natural settings, inside the bodies of the animals of which they are a part.

Recently, we developed a new in vivo model for studying collective cell migration (Bischoff *et al*., 2021). We examine the migration of the muscle precursors that will ensheathe and thereby sculpt the adult *Drosophila* testis in muscle. During pupal development, these cells move from the genital disc to the testis. There they migrate through a narrowly confined space between the pigment cells and the germline. They move collectively, loosely connected via N-cadherin-mediated cell-cell junctions, and their motility involves filopodial rather than lamellipodial protrusiveness (Bischoff *et al*., 2021). Our previous analysis revealed that these cells protrude and thus migrate toward the free cell edge of the sheet. This is a contact-dependent process, with Rho family GTPases regulating the stability of cell-extracellular matrix contacts, destabilizing those at cell-cell edges and stabilizing those at the free edge, thus promoting motility.

This system offers many advantages (Bischoff and Bogdan, 2021). The tissue can be cultured ex-vivo, with migration continuing for many hours. This feature, along with the very flat cell morphology, as cells squeeze between the pigment cell epithelium and the germline, allows live cell imaging at very high resolution, approaching that possible in cultured fibroblasts. In addition, the powerful genetic tools available in *Drosophila* allow precise manipulations. For example, one can drive gene expression specifically in the migrating myotubes using the GAL4-UAS system. Combining this with the virtually genome-wide library of UAS-drive RNAi reagents allows gene knockdown at will. It also allows expression of fluorescent proteins for visualizing cell protrusions, cell junctions or other subcellular structures. These tools helped power our previous work.

This system also allows us to explore a key issue in organogenesis: how one tissue shapes another. Post-migratory myotubes mature into circumferential muscles that shape the testis by an external “sculpting” process, reminiscent of mammalian lung morphogenesis, in which smooth muscles shape branching of the airway epithelium (Goodwin *et al*., 2019; Goodwin *et al*., 2023). After migration, myotubes elongate perpendicular to the proximo-distal axis, and in a convergent extension-like process, elongate the entire testis, resulting in its characteristic spiral shape (Rothenbusch-Fender *et al*., 2017; Bischoff and Bogdan, 2021). The external sculpting is fascinating, as it directly links collective migration to morphogenesis, but it is also helpful for performing a genetic screen, as it allows a quick read-out that reveals potential defects in migration and post-migratory shape changes. We can use adult testis shape to identify genes leading to defects in migration or post-migration morphogenesis. If the musculature covers only part of the testis, due to defects in migration, elongation and ultimately spiral-formation partially fails. This can cause the testis to have a dilated tip and the spiral to have fewer “revolutions”. If there is almost no coverage, testis retain their pupal ellipsoid shape. If the genital disc-seminal vesicle connection fails, myotubes cannot migrate at all, resulting in ellipsoid testes as well. Cell-cell adhesion defects or failure in active gap closure cause gaps in the muscle sheet, resulting in an uneven surface and kinks in the testis, also affecting the normal spiral shape (Rothenbusch-Fender *et al*., 2017).

Our current goal is to identify the proteins driving contact-regulated directed cell migration, and those involved in shaping the testis after migration. To identify these regulators, we took a two-step approach. First, we used RNA sequencing (RNAseq) to identify the “parts list” of the migrating cells—i.e., the genes expressed during migration.

We then used the literature and other resources to identify among these genes transmembrane receptors and transmembrane or secreted ligands that might be involved. To this list we added other genes known to regulate migration in other systems, and, because of the demonstrated roles of Rho-family GTPases, a set of regulators of these GTPases. We then obtained available RNAi reagents and used Mef-GAL4 to drive RNAi specifically in myotubes, knocking down genes one-by-one, and used the adult tests to identify hits. This revealed a long list of potential regulators. Together our gene expression data and RNAi screen provide the community with resources to further explore this fascinating process of collective cell migration and tissue morphogenesis

## Material & Methods

### Testis dissection and dissociation for RNAseq

We collected ∼100 timed U*AS-eGFP/+; dMef2/+* prepupae, either at 31 hrs APF or 45 hrs APF. We used 6 replicates per condition. We dissected pupal testes in ice cold M3 medium + BPYE (Shield and Sang) with 10% FCS (Thermo Fisher Scientific), and 1x Penicillin/Streptomycin (Gibco) and rinsed samples once in ice cold Cell Dissociation Buffer (Thermo Fisher Scientific). We dissociated testes in 500 µl ice cold Cell Dissociation Buffer with 2.5 mg/ml Collagenase (Invitrogen) and 4 mg/mg Elastase (Worthington Biochemical) in a glass block dish. Then we transferred the mix into a push cap tube and incubated it at 37°C in a water bath. During this step, we pipetted the mix up and down every 5 mins, using a p200 pipette, to effectively dissociate testes. We prefiltered cells using a 50 µm filter (Partec) and pelleted cells by centrifuging them 5 mins at 700 x g. We replaced the buffer with 1 ml ice cold FACS buffer (1x PBS, 2% BSA, 2.5 mM EDTA) and pelleted cells again by centrifuging them 5 mins at 700 x g. We replaced the buffer with 500 µl fresh ice cold FACS buffer. Then we added propidium iodide to a final concentration of 1 µg/ml and DyeCycle Violet to a final concentration of 5µM. We incubated samples 12 min at 37 °C in a water bath and FACS sorted them after that.

### FAC sorting and RNA isolation

FAC sorting was performed at the UNC Flow Cytometry Core Facility in Chapel Hill on a BD FACS Aria III. We sorted living (PI-negative) testis myotubes (GFP) that are bi- and tri-nucleated (DyeCycle Violet, distinct populations, see Figure 2). We sorted cells directly into 100 µl TRIzol LS and performed standard TRIzol RNA isolation by first adding 100 standard TRIzol and incubating the mix 15 mins at room temperature. We added 75 µl Chloroform, vortexed the mix for 30 seconds and incubated it for 3 mins at room temperature. We centrifuged the mix 20 mins at 4 °C and 12,000 x g. We transferred the aqueous phase to a fresh tube and added 180 µl Isopropanol and 1 µl Glycoblue. Then we vortexed the mix 15 seconds and precipitated the RNA over night at -80 °C. To pellet the RNA, we centrifuged the mix at 4 °C with 12,000 x g and washed the pellet twice with 75 % ethanol. We dried the pellet for 5 mins, to then resuspended it in 20 µl RNAse free water. RNAseq was performed at the UNC Advanced Analytics Core.

**Fig. 1.**
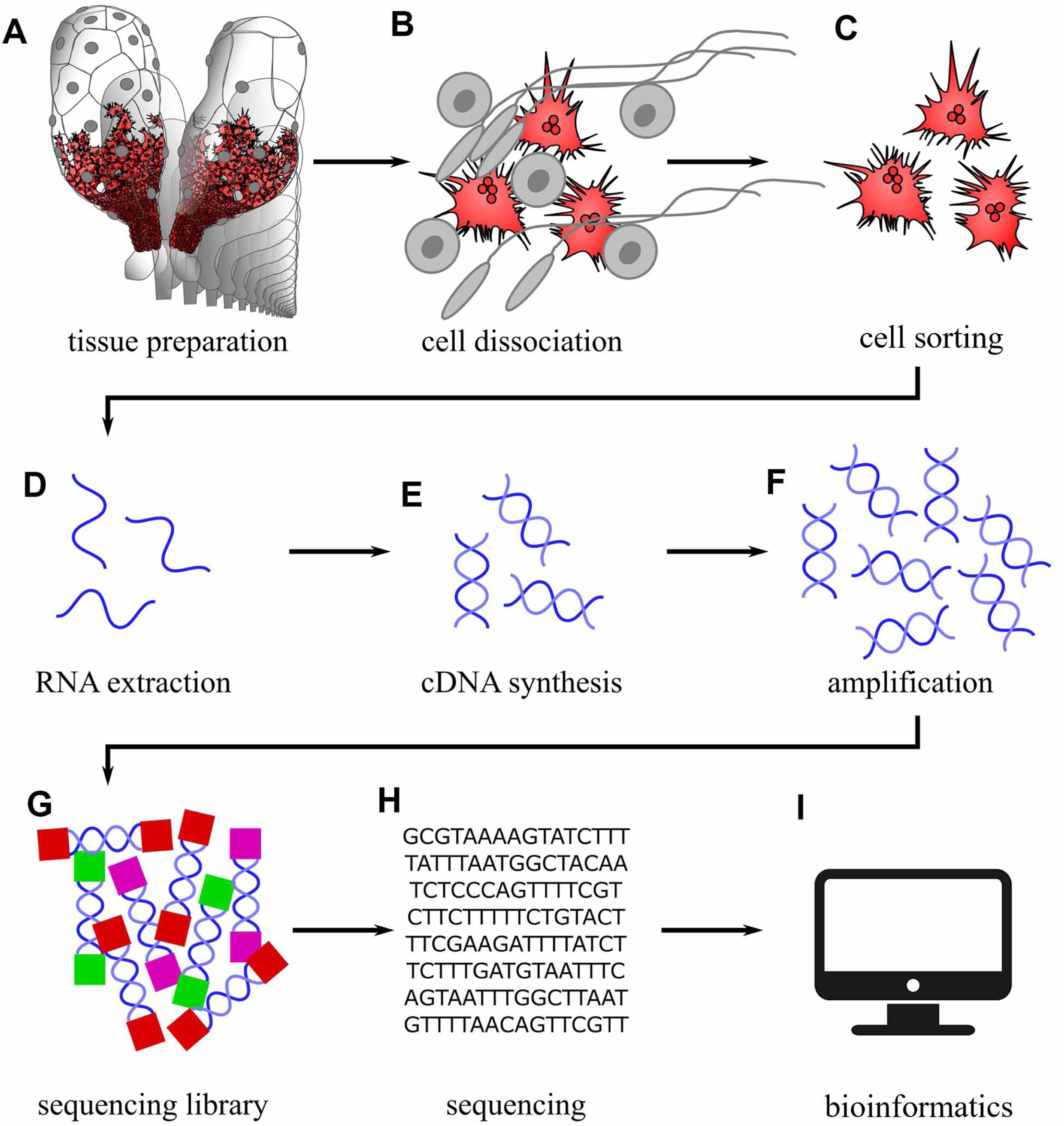
Diagram illustrating the steps used to generate our RNAseq datasets.

**Fig. 2.**
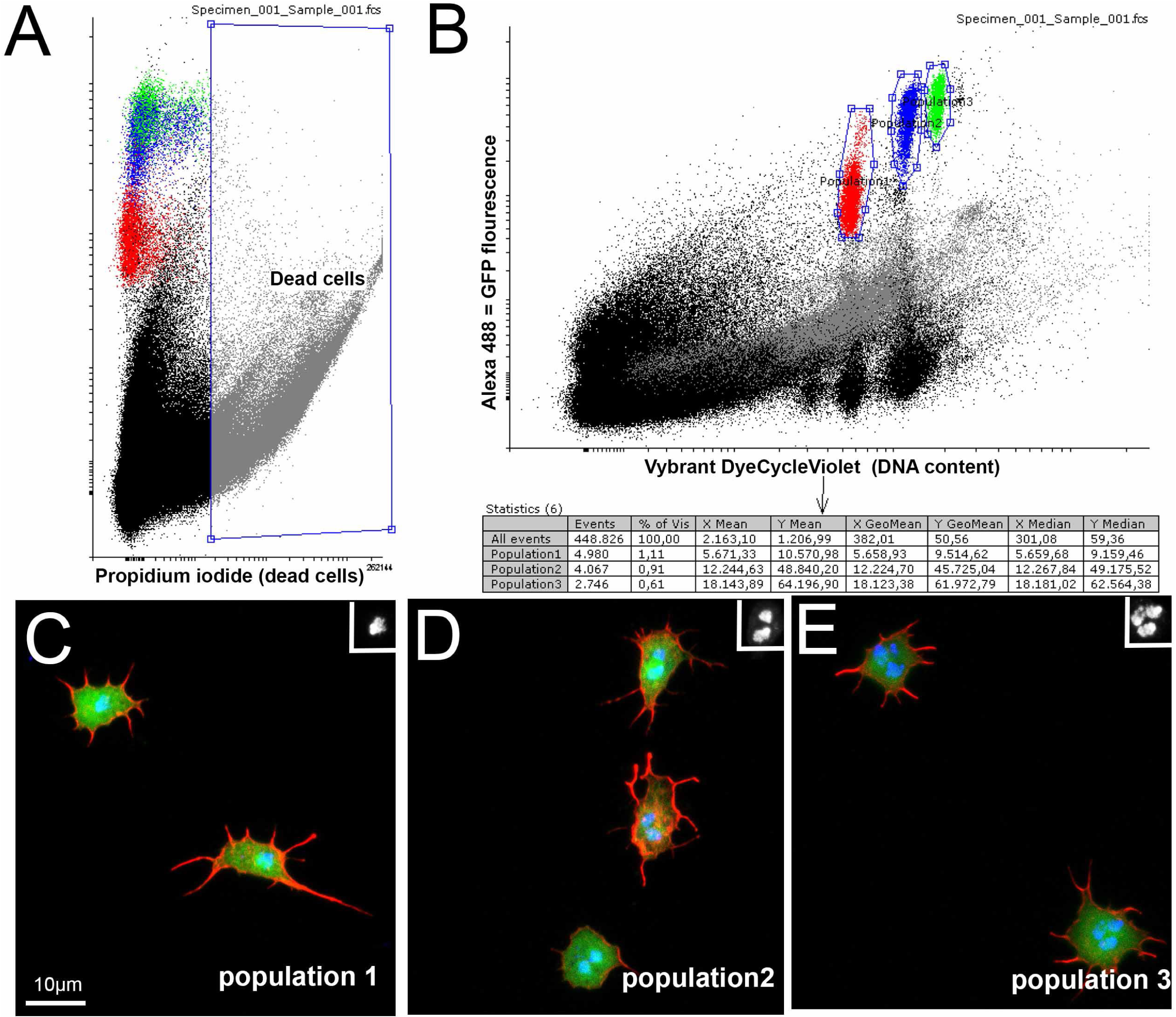
FACs sorting allowed us to isolate pure populations of living multinucleate myotubes. A. FACS profile along the propidium iodide versus the GFP-fluorescence [Alexa488] axis. Cells with high propidium iodide signals were likely dead cells and we discarded. B. FACS profile along the [v450] versus [Alexa488] axis. The Vybrant DyeCycleViolet [v450] signal assesses the DNA content of living cells and allowed us to separate the cells with high-GFP signal into three populations differing in DNA content. C-E. Examples of cells from the three populations revealing cells with one (C), two (D), or three (E) nuclei. Red=phalloidin (F-actin). Green=GFP Blue=DAPI (DNA). Insets in the upper right show the DNA channel of one cell from each image.

### Bioinformatic analysis

Quality control of the reads was performed with FASTQC (v 0.12.1). All runs passed metrics, with no poor quality reads. The Drosophila reference genome was downloaded from Flybase(dmel-all-aligned-r6.5, including the genome annotation file Drosophila_melanogaster .BDGP6.46.111.gtf). BBMap (v39.08) with default parameters was used to align the fastq files to the reference genome, which were then processed with Samtools (v1.21) to generate aligned .bam files. Read counts for gene models were determined using FeatureCounts (in the Subread v2.0.6) to produce a raw counts table from the .bam files. These counts were then used normalized by reads-per-kilobase-million RPKM). After normalization, RPKM estimates were used for visualizations and raw counts for downstream differential expression analysis using DESeq2 (v1.40.2).

The raw counts table was read into R (v4.3.1 including packages: data.table 1.15.4, dplyr 1.1.4, reshape2 1.4.4), and the non-protein-coding genes were filtered out. All further analysis was performed using R. DESeq2 compared the expression of the 6 replicate cell sets taken at 31 hours to 6 replicate cell sets taken at 45 hours. The model was *y ∼ timepoint + e.* The results were visualized in a principal component analysis plot using the ggplot2 graphing suite (ggplot2 3.5.1). Gene tables for several classes of functionality were cross-referenced with the results to produce boxplots illustrating the differential expression levels of the key genes. Three-dimensional volcano plots were produced using the Glimma package (v2.12.0) and a dynamic instantiation is hosted here: https://rpubs.com/Cserody/bischoff_volcano_symbols. An FGSEA analysis was performed using the FGSEA package for R (fgsea1.28.0, also EnrichmentBrowser 2.32.0, ggrepel 0.9.5, and dynamicTreeCut 1.63.1), and its native writeGMT and getGenesets functions were used to generate the gmt and genesets files. For FGSEA, we limited to GO terms with over a 10 and under 1000 members.

### Fly genetics

We performed crosses at 25 °C. Prepupae for subsequent live-cell imaging developed at 26.5 °C in a cell incubator. Fly stocks used in the screen can be found in Supplemental Table 5. For control crosses, we used *w^1118^* and referred to it as wildtype or WT.

### Testis staining and microscopy

For the screen, testes from adult flies were dissected 1–3 days post-hatching in 1.5x PBS and fixed 20 mins in 4% PFA in PBS. Subsequently, we washed them three times in 1.5x PBS and once in 1.5x PBS with 0.1% Tween 20. Overnight phalloidin stainings were performed using 1:500 Phalloidin (*ThermoFisher, Alexa Fluor 488*) in 1.5x PBS with 0.1% Tween 20 at 4°C. After washing samples three times using 1.5x PBS, we mounted them in PBS in 35 mm glass bottom dishes. For imaging we used an LSM Pascal (Zeiss) with a 10x (Zeiss EC Plan-Neofluar 10x/0.3) dry objective. All images of adult testes are projections of large z-stacks using maximum intensity. We modified brightness and contrast in Photoshop (Adobe) and Fiji (ImageJ) to make all structures visible.

### Life cell imaging

We dissected testes from 30 h APF testes in M3 Medium (Shields and Sang, Sigma Aldrich) with 10% FCS (Thermo Fisher Scientific), and 1x penicillin/streptomycin (Gibco) at room temperature (Bischoff and Bogdan, 2023). For mounting them live, we used 0.5 % low gelling agarose (Sigma Aldrich) in M3 Medium. For image acquisition we used a Nikon Ti2 inverted microscope with a Yokogawa CSU-W1 spinning disk and Hamamatsu ORCA-fusion BT sCMOS camera and a 25x/1.05 Silicone Apochromat objective. Using Fiji (ImageJ), we changed *Gamma* to 0.5, to make all parts of the tissue visible without oversaturating other parts of the image. We modified brightness and contrast in Photoshop (Adobe) to make all structures visible.

### Cdc42/Rac activity sensor quantification

To quantify Cdc42/Rac activity, we used a characterized sensor that has the CRIB domain of the *Drosophila* Pak2 gene *mbt* fused to GFP and under control of a UAS-promoter (Rötte *et al*., 2024). To quantify its effects, we expressed it in testis myotubes using the *dMef2-Gal4* driver. As it is naturally strongly enriched at cell-cell borders, we used a line scan-method to quantify it in wildtype or after expressing RNAi constructs. To that end, we wrote a Fiji script. The user must mark cell-cell borders and the script generates a line of defined length perpendicular to that line, with the center exactly on top of the cell-cell border so that multiple profile measurement can later be overlayed. We averaged the measurements within each sample and then normalized them by setting the minimum of the averaged values to 0. To simply show the fluorescence intensity at the membrane in one dimension, we used Fiji to create lines on top of cell-cell contacts and measured the averaged fluorescence intensity.

### Detailed author contributions

Maik Bischoff conceived the project, prepared the samples for RNAseq, analyzed the outcome to determine which genes to include in the screen, directed the team who carried out the screen, and placed the genes in phenotypic categories. Jenevieve Norton carried out much of the screen, organized the data, and trained undergraduates who participated in the screen, Erika Munguia also made substantial contributions to the screen, assisted by the team of Sarah E. Clark, Rebecca Korankye, Emmanuel Addai Gyabaah, Taino Encarnacion. Noah Gurley helped prepare samples for RNAseq. Christopher Serody and Corbin Jones did the bioinformatics analysis. Maik Bischoff, Corbin Jones and Mark Peifer wrote the paper with editorial contributions from the other authors.

## Results

### Identifying genes expressing in migratory-stage myotubes using RNAseq

The first step in our effort to identify regulators of myotube collective cell migration was to identify candidate genes expressed in these cells. To do so, we used RNA seq to identify both genes expressed in migratory stage nascent myotubes, and those expressed in myotubes after migration was completed. To identify myotubes in this complex tissue, we used flies in which the muscle specific *dMef2-Gal4* driver drives *UAS-eGFP*. We dissected six sets of 150-200 testis from pupae 31 hr after pupae formation (APF), a timepoint in the middle of the migratory phase, or six sets of 150-200 testis from pupae 45 hr APF, a timepoint after the completion of migration (Figure 1A). We disassociated cells using Collagenase and Elastase (Fig 1B; (Rust *et al*., 2020). To identify the right population of cells for sorting, we stained them with propidium iodide (PI), which stains DNA in dead cells but is not taken up by living cells, and DyeCycle violet, a cell permeable DNA dye, and sorted them by FACS (Fig 1C).

We discarded cells with high PI staining (Fig. 2A), as they represented dead cells. When analyzing levels of eGFP and DyeCycle violet, we could easily identify three populations of TNM with high GFP-expression, revealing them to be myotubes of the genital tract. On the DyeCycleViolet axis, myotubes formed three distinct populations (1-3) – consistent with differing numbers of nuclei (Figure 2B). Microscopic analysis revealed population 1 to be mononucleated (Figure 2C). The only mononucleated cells in the dissected tissues are myotubes of the ejaculatory duct and the paragonia (Susic-Jung *et al*., 2012). Micrographs of population 2 and 3 revealed them to have 2 or 3 nuclei, respectively, thus clarifying that these are the multinucleated myotubes that migrate on the testis (Figure 2D,E). Therefore, we combined population 2 and 3 for the subsequent library preparation.

We next used a TRIzol protocol to obtain RNA from FAC-sorted myotube populations 2 and 3 (Fig. 1D). Subsequently, we performed Qbit and Bioanalyzer quality control. cDNA was made and amplified (Fig. 1E,F). We used the Takara SMART-Seq v4 Kit for library preparation, that included Poly-A selection (Fig 1G). For sequencing we used the NextSeq P3 flow cell method with a 2 x 50 bp paired end format (Fig 1H). We worked with mean-values of paired ends and lanes. Principal component analysis showed that the 31 hr condition and the 45 hr condition each cluster, with 92% variance between them and only 3 % variance within conditions (Suppl. Figure 1). For quality control, we used FastQC. FASTQC showed that most replicate had high depth of coverage with a minor exception of one replicate form 45h that had lower reads (average number of reads for 31h: 109.7M; St. deviation 14.5M; average number of reads for 45h: 89.0M; St.Dev. 28.7M). All replicates had high quality reads and no problematic reads.

These data provided us with the parts-list of migrating myotubes (Suppl. Table 1). We took two approaches to identify potential candidates from this list, looking for genes that were differentially upregulated in migrating myotubes, and also looking for candidate ligands, receptors and potential downstream signaling proteins that were expressed in migrating myotubes, even if they were not substantially downregulated in the later timepoint.

We normalized the data using both FPKM and DESeq2 (v1.40.2) estimates of dispersion parameters both sample- and gene-wise (Suppl. Table 2). We performed gene Set enrichment analysis (FGSEA, version 1.28.0) using the genes showing strong differential expression as a “biological quality control”. Consistent with our expectations, the gene ontology (GO) terms “*myofibrill assembly”*, “*muscle attachment”* and “*sarcomere organization”* were among the highest upregulated during the 45 hr time step – confirming the validity of the data set (Fig. 3A; Suppl. Table 3). Gene ontology terms related to synaptic development were also enriched, perhaps reflecting the assembly of neuromuscular junctions.

**Fig. 3.**
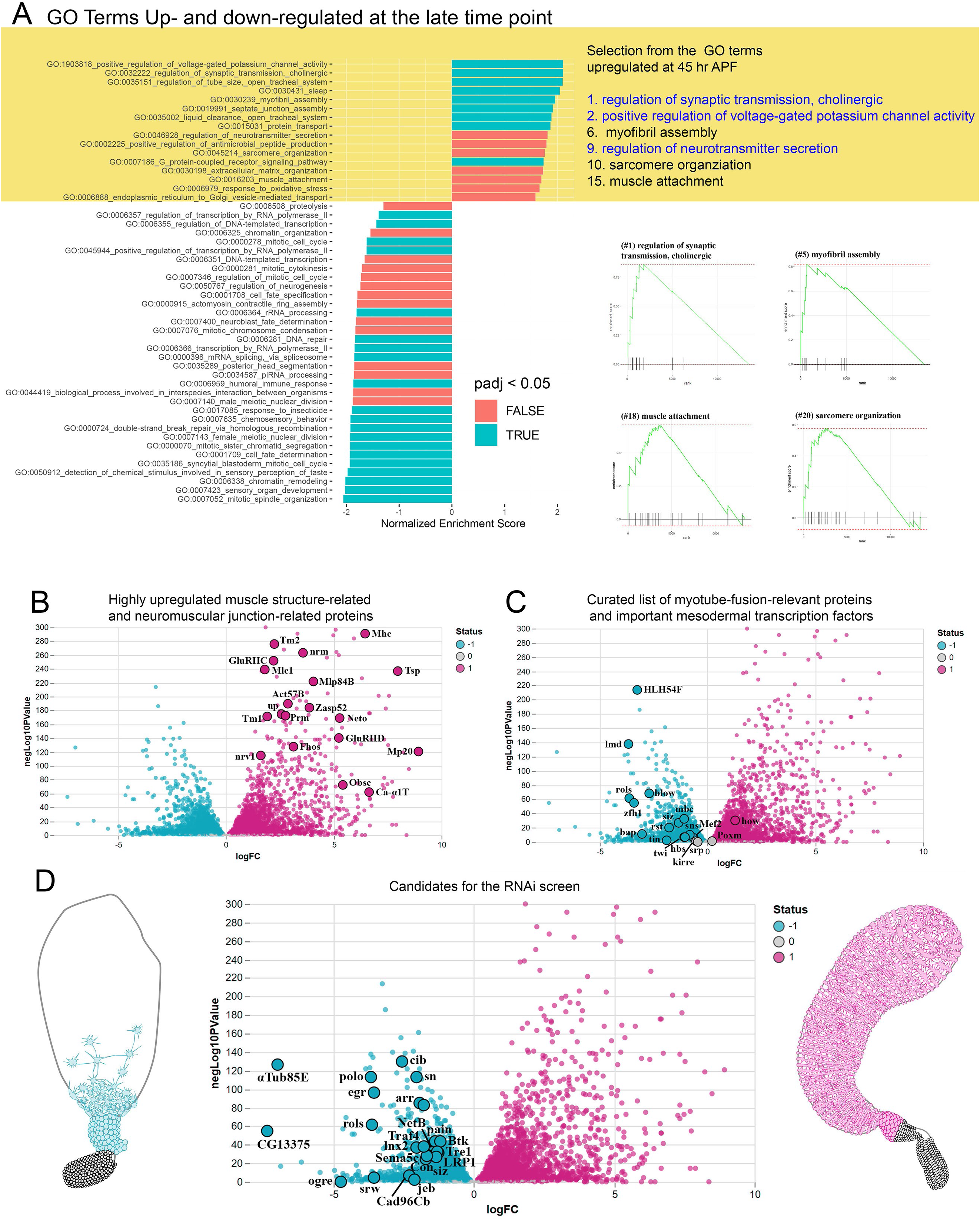
Identifying genes up- and down-regulated at the early and late timepoints. A. Gene ontology (GO) terms of genes up-regulated (top, highlighted in gold) and down-regulated (bottom) at th 45 hr APF timepoint. At the right are examples of up-regulated GO terms that reflect muscle development or potential neuromuscular synapse assembly. B-D. Volcano plot of genes upregulated at the 30 hr APF (left, highlighted in teal) or the 45 hr APF timepoint (right, highlighted in magenta). B. Highlighted are selected examples of genes with known roles in muscles that are upregulated at the 45 hr APF timepoint. C. Highlighted are selected examples of genes that encode mesoderm transcription factors or proteins involved in myotube fusion that are upregulated at the 30 hr APF timepoint. D. Highlighted are 22 genes that are upregulated at the 30 hr APF timepoint which we chose to include in the RNAi screen.

Next we inspected the DeSeq2 data manually, and this revealed that many of the most upregulated genes in the post-migratory stage (45 hrs APF) are directly related to sarcomere and neuromuscular junction formation, such as the muscle myosin heavy and light chains (Mhc and Mlc1), Tropomyosin (Tm2), and Neuromusculin (nrm), a protein involved in synaptic target recognition (Fig. 3B). We also created an interactive version of the Volcano plot for others who want to analyze the data looking for different features (https://rpubs.com/Cserody/bischoff_volcano_symbols). During the migratory stage, we expected a higher abundance of early mesodermal transcription factors and of genes that regulate myotube fusion, which occurs directly before migration. Since these GO-terms did not exist, we analyzed the DeSeq2 data manually for typical genes with these functions. Indeed, 13 out of 14 selected genes are relatively upregulated during the migratory time-step, including Lame duck (*<u>lmd</u>*), a zinc finger transcription factor essential for the specification of fusion competent myotubes and for myotube fusion (Duan *et al*., 2001), HLH54F, an HLH transcription factor required for the specification and migration of longitudinal gut muscle founders (Ismat *et al*., 2010), and Rolling pebbles (*rols*), a protein required for myotube fusion in founder cells (Fig. 3C; 31 hrs APF; (Menon and Chia, 2001)), further validating the data. The full list of differentially expressed genes is in Suppl. Table 2. These data will provide a resource for all wanting to explore the molecular mechanisms underlying myotube migration or testis morphogenesis.

### Selecting candidate genes for our RNAi screen

We then used the 31 hr APF gene list to identify potential regulators of migration. Our first list of candidate genes was derived by manually examining the genes that were upregulated during the 31 hr time step (Figure 3D). From these we chose a list of 22 genes. To identify additional candidates among genes that were expressed during migration and did not get downregulated, we took a different approach. We first identified sets of proteins of interest—e.g. transmembrane receptors--using the Flybase-curated “*Gene Group”*-lists. We analyzed multiple gene groups in that way, normalized the count data to the total number of reads and to gene length (reads per kilobase of transcript per million reads mapped, RPKM; Suppl. Table 4) to look for highly expressed members of each group (Fig. 4).

**Fig. 4.**
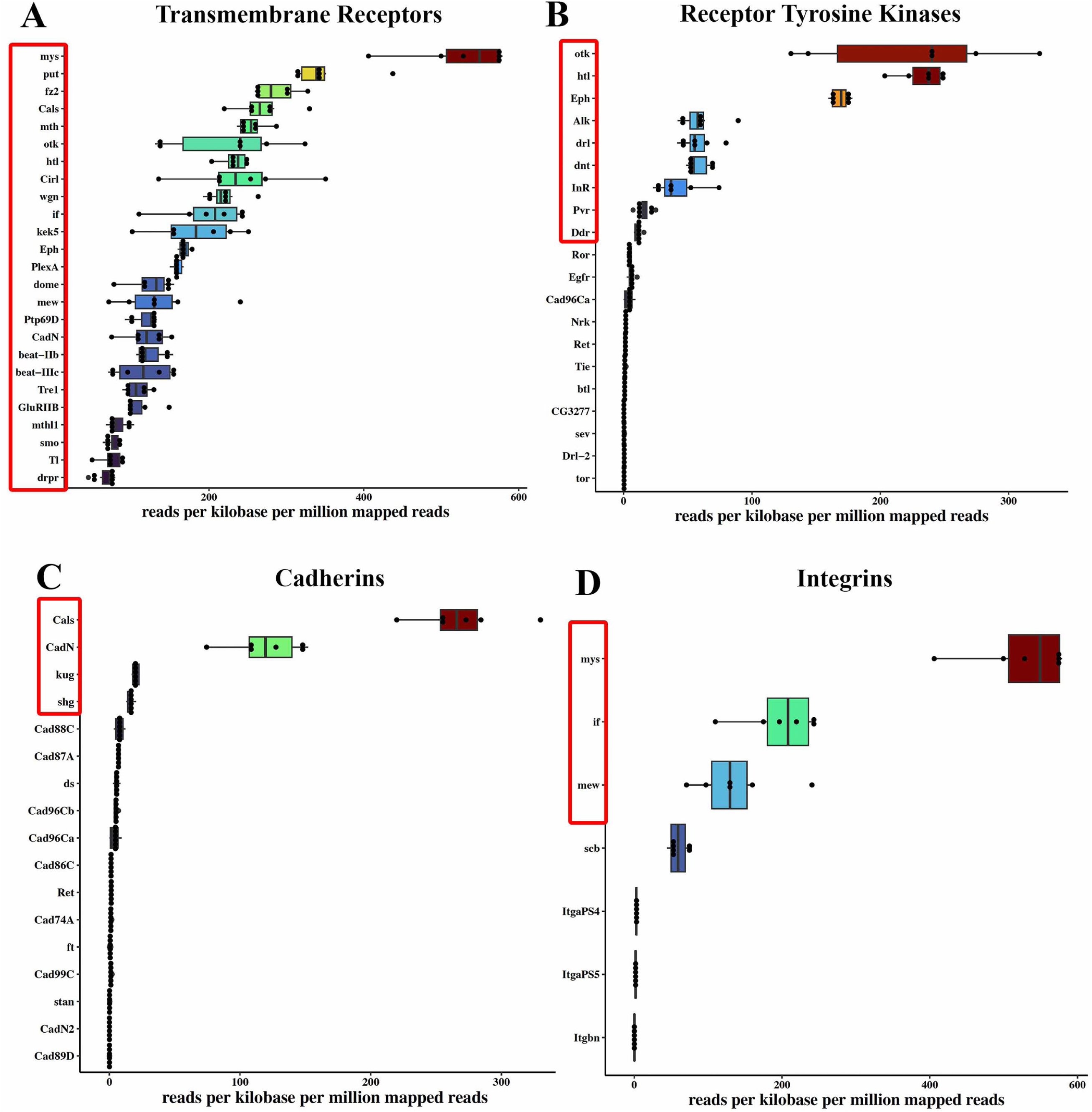
Genes selected for our RNAi screen. Four lists derived from the Flybase-curated “Gene Group”-lists. In each genes are ranked by level of expression (reads per kilobase per million mapped reads), and genes included in our RNAi screen are boxed in red. A. Transmembrane receptors. B. Receptor tyrosine kinases. C. Cadherin family members. D. Integrin subunits.

Since we were interested in directed cell migration, we focused on transmembrane receptors. Therefore, we focused on the 25 most highly expressed genes in the “*transmembrane receptor*” gene group (Fig. 4A). Two genes that were previously known to be functionally important for TNM migration were highly expressed, including the FGF receptor *heartless* (*htl*) (Rothenbusch-Fender *et al*., 2017), and the Integrin betaPS subunit *myospheroid* (*mys*) and thus we excluded *htl* from our screen. We supplemented our list with the most highly expressed receptor tyrosine kinases (RTKs) that were not already within the top 25 transmembrane receptors but were still among the 9 highest expressed RTKs (Fig. 4B). Interestingly, multiple transmembrane receptors that are involved in axonal pathfinding – a process related to contact-dependent cell migration – are highly expressed during migration (31 hr APF, Figure 3D). These include the classical axon guidance factors *Plexin A* (*PlexA*), the Plexin coreceptor *off-track* (*otk*), the Plexin B ligand *Semaphorin 2a* (*Sema2a*), *Netrin B* (*NetB*), *beaten path IIb* (*beatIIb*), *beaten path IIIc* (*beatIIIc*), the Ephrin receptor tyrosine kinase (Eph) and the Latrophilin homologue *Calcium-independent receptor for α-latrotoxin* (*Cirl*) (Figure 4A,B).

In parallel with identifying potential transmembrane receptors, we also looked for possible transmembrane or secreted ligands that might be involved. From the “*receptor ligand*” gene group, we included the three most highly expressed genes, *mav*, *miple1* and *Semaphorin2a* in our candidate list. Since Wnt-signaling is known to play a role in TNM development (Kozopas *et al*., 1998; Rothenbusch-Fender *et al*., 2017), we added to two most highly expressed Wnt-genes, *Wnt4* and *Wnt5*.

Cell-cell and cell-matrix adhesion also play important roles in cell migration. As signals might be transduced directly via cadherin family receptors, we analyzed the expression of all Cadherins (Fig. 4C). Consistent with our previous finding that *N-Cadherin (Cad-N)* plays important role in TNM migration (Bischoff *et al*., 2021), it was the second most highly expressed cadherin. We thus included *Cals*, *kug,* and *shg* on our candidate gene list. We also included the three most highly expressed integrin subunits (Fig. 4D). Since proteolytic degradation of the ECM environment might be involved in TNM migration we opted to include the highly expressed Metzincin matrix metalloproteases *AdamTS-A*, *kuz* and *AdamTS-B*.

In addition to proteins involved in cell-cell signaling, we also explored a subset of cytoskeletal regulators. Pharmacological perturbations revealed that Formins play an important role in TNM migration (Bischoff *et al*., 2021). We thus added the highest expressed Formins DAAM and Frl to our list, and also added the actin-regulatory F-Bar protein Cip4. Finally, Rho family GTPases are fundamental for TNM migration (Bischoff *et al*., 2021). Guanine nucleotide exchange factors (GEFs) activate these GTPases by stimulating the release of GDP to allow GTP binding, while GTPase-activating proteins (GAPs) turn GTPases off by binding to activated G proteins and stimulating their GTPase activity. We thus included all existing Rho-family GEFs and GAPs to the screen. Due to their important role in the regulation of cell adhesion, we also included all Flybase-annotated Rap1 GTPase GAPs and GEFs.

### The phenotypes observed in our screen reveal the complexity of events that shape this organ

Using this gene set, we examined the collections of UAS-RNAi reagents at the Vienna and Bloomington *Drosophila* Stock centers, and ordered one or more RNAi lines for each gene. Transgenes regulated by the UAS sequences can be activated the GAL4 transcription factor. We used a line in which GAL4 is under control of the promotor of the muscle-specific gene *Mef4.* This allowed us to knockdown gene expression in a tissue-specific manner, and assess the potential role of each gene in myotube migration. Females carrying *Mef2-GAL4* were crossed to males carrying each UAS-RNAi line. One-to-three-day old adult male progeny were collected and testis dissected from 10 males, replicated three times. Testis were examined under a dissecting scope. For any lines with potential morphological defects, testes were then stained with fluorescently labeled phalloidin, which binds F-actin and thus outlines muscles. These were then imaged by confocal microscopy.

We tested 279 RNAi lines covering 131 genes. A subset of the RNAi lines led to embryonic or larval lethality, precluding analysis of adult phenotypes (Suppl. Table 5). Some led to pharate adult lethality, before adult eclosion—for these genes, testes were dissected from pharate adults. We identified morphological defects for 29 genes (Table 1). These varied widely in the strength and nature of the defects observed.

Our original goal was to define the factors required for testis nascent myotube migration. However, as our screen proceeded and phenotypes began to emerge, we realized our mutant collection included genes that are required for diverse processes in testis morphogenesis. The wildtype testis is full covered in circumferential muscle, has a spiral shape with a very gradual reduction in diameter toward the proximal end (Fig. 5A). Some knockdowns had phenotypes resembling those we had anticipated from defects in migration. In these there was a failure to cover the testis in muscle, ranging from weaker effects when on the distal tip was uncovered to more severe effects with substantial loss of muscle coverage. The two strongest effects were seen when knocking down *RtGEF*/*dPix* (Fig. 5B), a Rho-family GEF targeting Rac1 and Cdc42, the matrix-metalloprotease Kuz (Fig. 5C; one of the two RNAi lines), and the Jak/Stat receptor Dome (Fig. 5D; one of the two RNAi lines). For these genes, knockdown led to dramatic testis shape changes, with a highly expanded distal end and shortened proximal distal axis. The line targeting Dome also had an interesting additional effect: the muscles remaining appeared striated in morphology (Fig. 5D, inset), suggesting defects in cell fate determination.

**Fig. 5.**
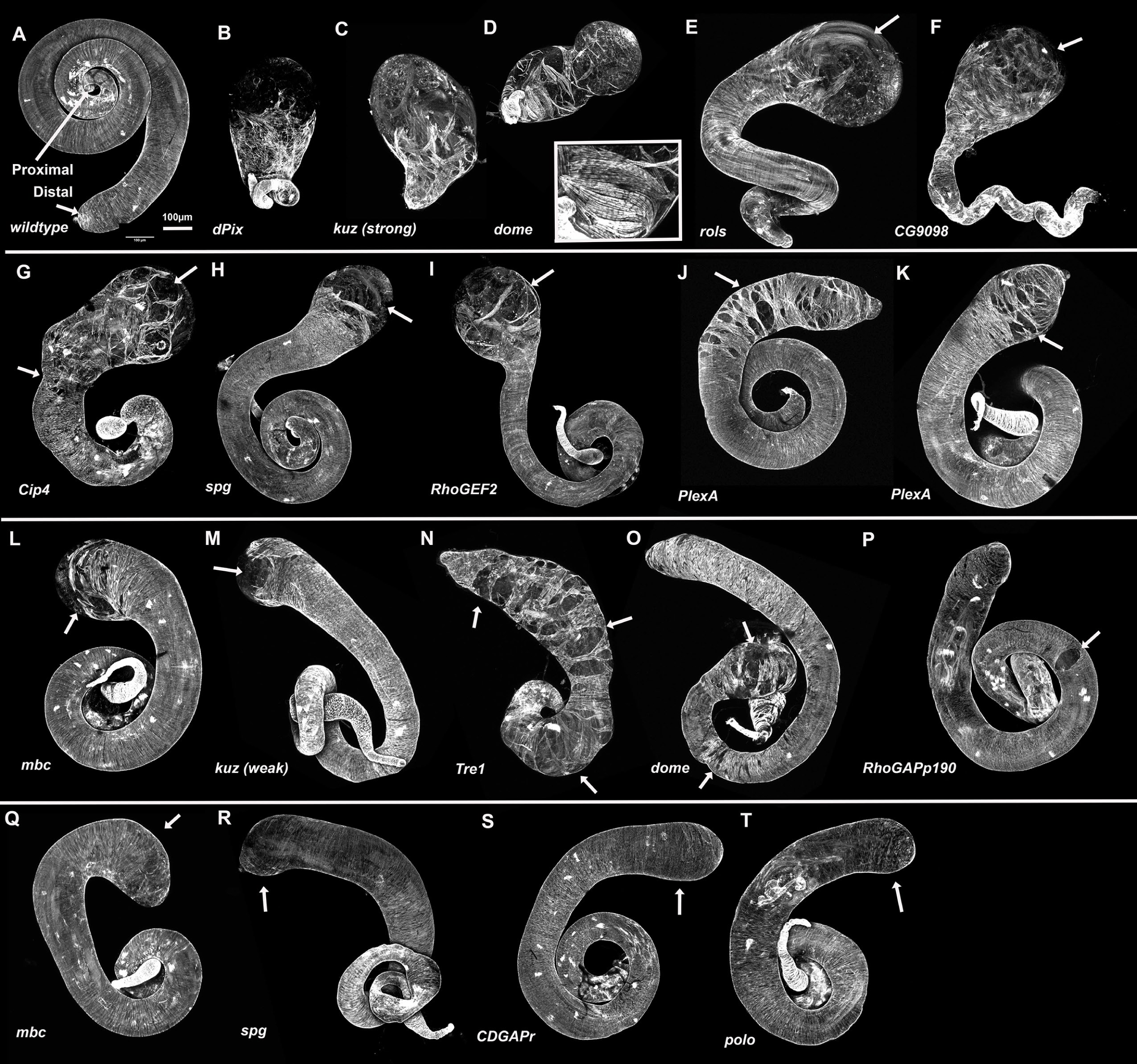
RNAi lines that alter muscle coverage or distal testis shape. Adult testes from wildtype or adults expressing a UAS-driven RNAi line targeting the indicated gene under control of *mef-*GAL4. All are stained with fluorescently-labeled phalloidin to reveal F-actin, which highlights muscles. A. Wildtype. The wildtype testis is fully covered in circumferential muscle and has a spiral shape with a very gradual reduction in diameter toward the proximal end. B-T. Testis from knockdown lines, arranged according to phenotypic class. Detailed descriptions of each are in the text. B-D. Extreme loss of muscle coverage. The inset in D shows a close-up illustrating the “straited muscle phenotype”, E-I. Strong loss of distal muscle coverage (arrows). The more proximal arrow in G shows variation in testis diameter. J-M. Variable loss of distal muscles or gaps in muscle coverage (arrows). N-P. Strong (N) or weaker gaps that are not confined to the distal end (arrows). Q-T. Distal testis is enlarged (arrows).

The next set of genes included those for which RNAi led to strong, moderate or mild loss of <u>distal</u> muscle coverage. Very strong loss of distal muscle coverage was seen after knockdown of the adapter protein Rols (Fig. 5E), CG9098, an uncharacterized predicted GEF with an N-terminal SH2 domain whose Nsp family mammalian orthologs are predicted to lack enzymatic activity and instead act as adapters (Fig. 5F), and the F-Bar protein Cip4 (Fig. 5G). CG9098 knockdown also led to narrowing of the proximal testis, while Cip4 knockdown also led to additional alterations in overall testis shape. Knockdown of the DOCK family RhoGEF Spg (Fig. 5H; one of two lines), RhoGEF2 (one of two lines; Fig. 5I), and the axon guidance receptor PlexA (two lines; Fig 5J, K) all caused moderate loss of distal muscle coverage. More mild loss of distal muscle coverage was seen after knockdown of the unconventional bipartite Rac-GEF Mbc (Fig. 5L; one of two lines) and the matrix-metalloprotease Kuz (Fig. 5M; one of two lines).

Some RNAi lines led to gaps all along the sheet, perhaps reflecting milder defects in migration, or defects in cell adhesion or the ability to actively close gaps. Knockdown of the G-protein coupled receptor (GPCR) Tre1 led to extensive muscle coverage gaps that extended much more proximally (Fig. 5N). One of the RNAi lines knocking down the Jak/STAT receptor Dome has similar but much weaker gaps restricted to the proximal part of the testis (Fig. 5O), a phenotype also caused by knockdown of *RhoGAPp190D* (Fig. 5P).

However, these were only a subset of the RNAi lines that altered the adult testis. More surprising to us were mutants in which there was nearly complete or full muscle coverage, but in which the final muscle sheet seemed to have different physical properties, not allowing normal tissue shaping. Some of these had relatively mild effects. One of the lines knocking down Mbc lead to broadening of the distal third of the testis (Fig 5Q), while one of the lines knocking down the DOCK family RhoGEF Spg (Fig. 5R), the line knocking down the Rho family GAP CDGAPr (Fig. 5S) and one line knocking down the cell cycle kinase Polo (Fig. 5T) led to mild broadening of the distal tip—many of the testes affected by this *polo* RNAi line failed to connect to the seminal vesicle and thus muscle migration never started.

Others affected the “shaping” of the testis in different ways. Some lines affected the consistency of the testis diameter, with a resulting “waviness” of the testis border. Three RNAi lines led to both broadening of the distal end and a waviness to the testis border due to variations in diameter. These included knockdown of the axon guidance ligand NetB (Fig. 6B) and two lines knocking down the alpha-tubulin isoform atub85E (Fig. 6C, D). Knockdown of the Ras/Rap family GAP Gig or the BTB/POZ domain protein Rsh led to variation in testis diameter without distal enlargement (Fig. 6E, F). Knockdown of the RhoGAP Rlip and one of the lines targeting RhoGEF2 broadened and slightly shortened the whole testis, with (Rlip) or without variation in testis diameter (Fig. 6G, H). Knockdown of the RhoGEF Pura had a very curious effect—broadening the distal tests while narrowing the proximal testis (Fig. 6I). In contrast, two of the lines targeting the Ras GEF Sos led to narrowing only of the distal testis tip (Fig. 6J, K). Finally, knockdown of the FERM domain-containing protein Cdep or RhoGEF64C narrowed and elongated the entire testis (Fig. 6L, M).

**Fig. 6.**
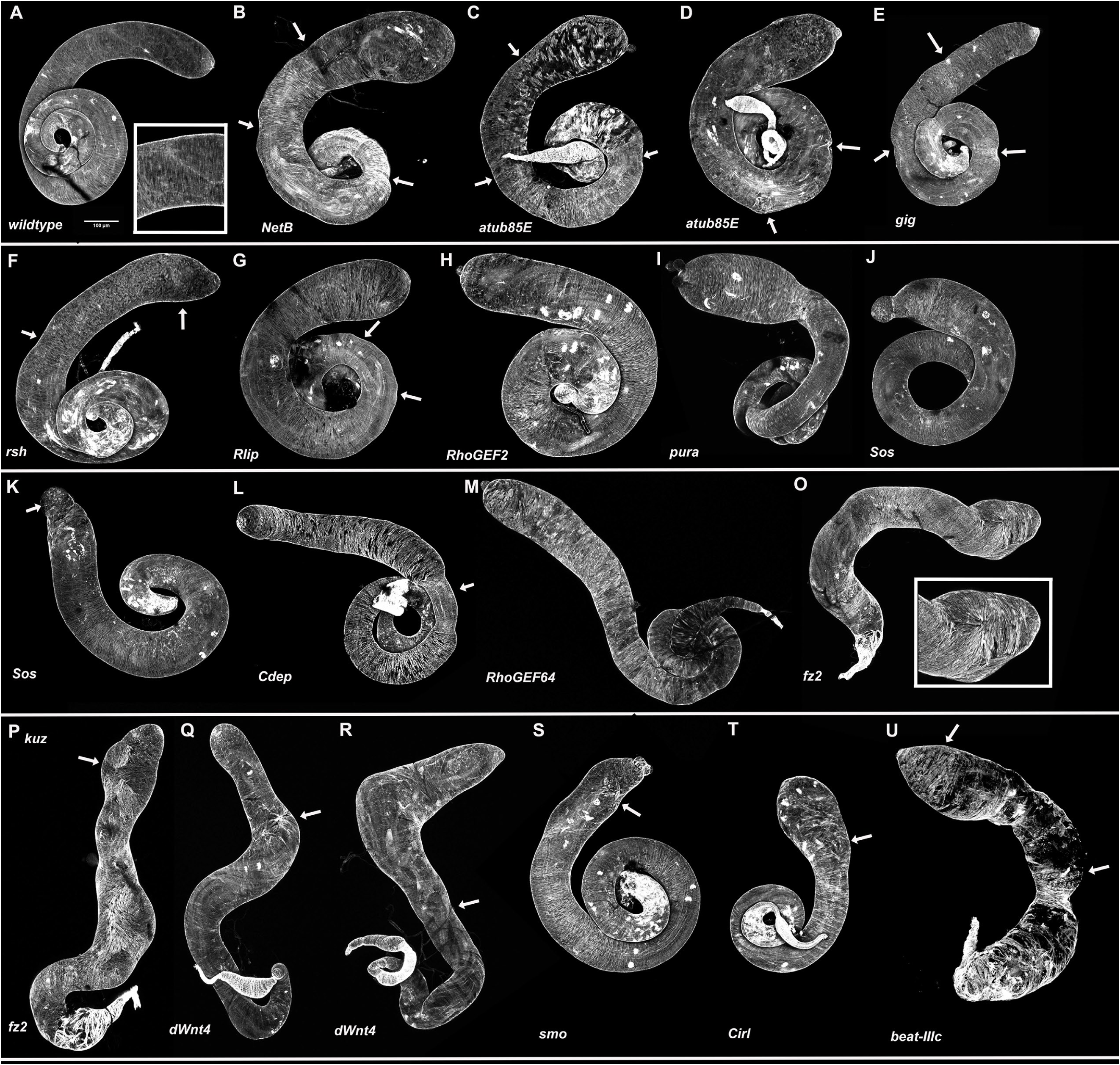
RNAi lines that alter testis morphology in other ways. Adult testes from wildtype or adults expressing a UAS-driven RNAi line targeting the indicated gene under control of *mef-*GAL4. All are stained with fluorescently-labeled phalloidin to reveal F-actin, which highlights muscles. A. Wildtype, showing normal shape. Inset reveals normal aligned muscles. B-F. Variable testis diameter leading to a wavy margin (arrows). G-H. Broadened and shortened testis. Rlip knockdown also leads to variations in diameter (arrows). I-L. regional variations in testis diameter. M. Narrowed and elongated testis. O-R. Strong defects in muscle alignment (Inset in O, arrows) and loss of spiraling. S-U. Defects in muscle alignment (arrows) coupled with other defects in testis shape.

In wildtype the muscles are well aligned into a circumferential pattern (Fig. 6A, inset). A subset of the RNAi lines altered this normal parallel circumferential alignment, perhaps reflecting problems in the parallel alignment/intercalation phase. In the testis of these lines spiral formation completely fails. Two of the genes with this phenotype form a ligand/receptor pair: two RNAi lines targeting the gene encoding the Wnt receptor *fz2* (Fig. 6O, inset, P) and two lines encoding the ligand *dWnt4* (Fig. 6Q, R). The RNAi line targeting gene encoding the Hedgehog receptor *Smo* altered muscle arrangement and testis shaping distally (Fig. 6S) while an RNAi line targeting the G-protein coupled receptor *Cirl* led to a somewhat similar phenotype, with the distal end of the testis enlarged and some alterations in muscle arrangement (Fig. 6T). Knockdown of the Ig-family receptor Beat-IIIc led to a complex phenotype including failure of testis spiraling, extremely variable testis diameter and muscle alignment defects (Fig. 6U). Intriguingly, over-expression of Beat-IIIc and loss-of-function of Wnt4 and Fz2 all altered motor neuron synaptic specificity (Inaki *et al*., 2007)

Finally, a few other RNAi lines had more complex effects. Several RNAi lines led to apparent failure of the genital disc to connect to the testis in some individuals: these included lines targeting the genes encoding *polo*, *dPix*, and *rols* (Suppl. Table 1). Finally, in animals in which the RNAi line targeted the gene encoding *RapGAP1*, the testis was extremely small (Fig. S1A vs B), though the remnant was covered in muscle (Fig. S1C), suggesting a possible non-tissue autonomous effect.

### Beta-pix RNAi leads to very strong defects in cell migration

To illustrate the utility of these combined RNAseq and RNAi datasets, we examined one of our hits further. Among the strongest phenotypes observed in our screen was that caused by knockdown of dPix (also known as *RtGEF),* the Drosophila homolog of the mammalian Rho-type guanine nucleotide exchange factor beta-PIX (Zhou *et al*., 2016). After *dPix^RNAi^*, many testes were still round and largely uncovered in muscle, and in less severe examples the testis was shortened in the proximal distal axis and the distal end was uncovered (Fig. 7A,B). dPix has known roles in synaptic structure and growth in the nervous system (Parnas *et al*., 2001; Ho and Treisman, 2020), in maintaining epithelial architecture and collective cell migration of the follicle cells during oogenesis (Dent *et al*., 2019), and in regulating Hippo signaling in imaginal discs (Dent *et al*., 2015). The RNAi line we used is a well validated one, which is known to reduce *dPix* mRNA, and to mimic the effect of the strong *dPix^1036^* mutant in effects on Hippo signaling in imaginal discs (Dent *et al*., 2015).

**Fig. 7.**
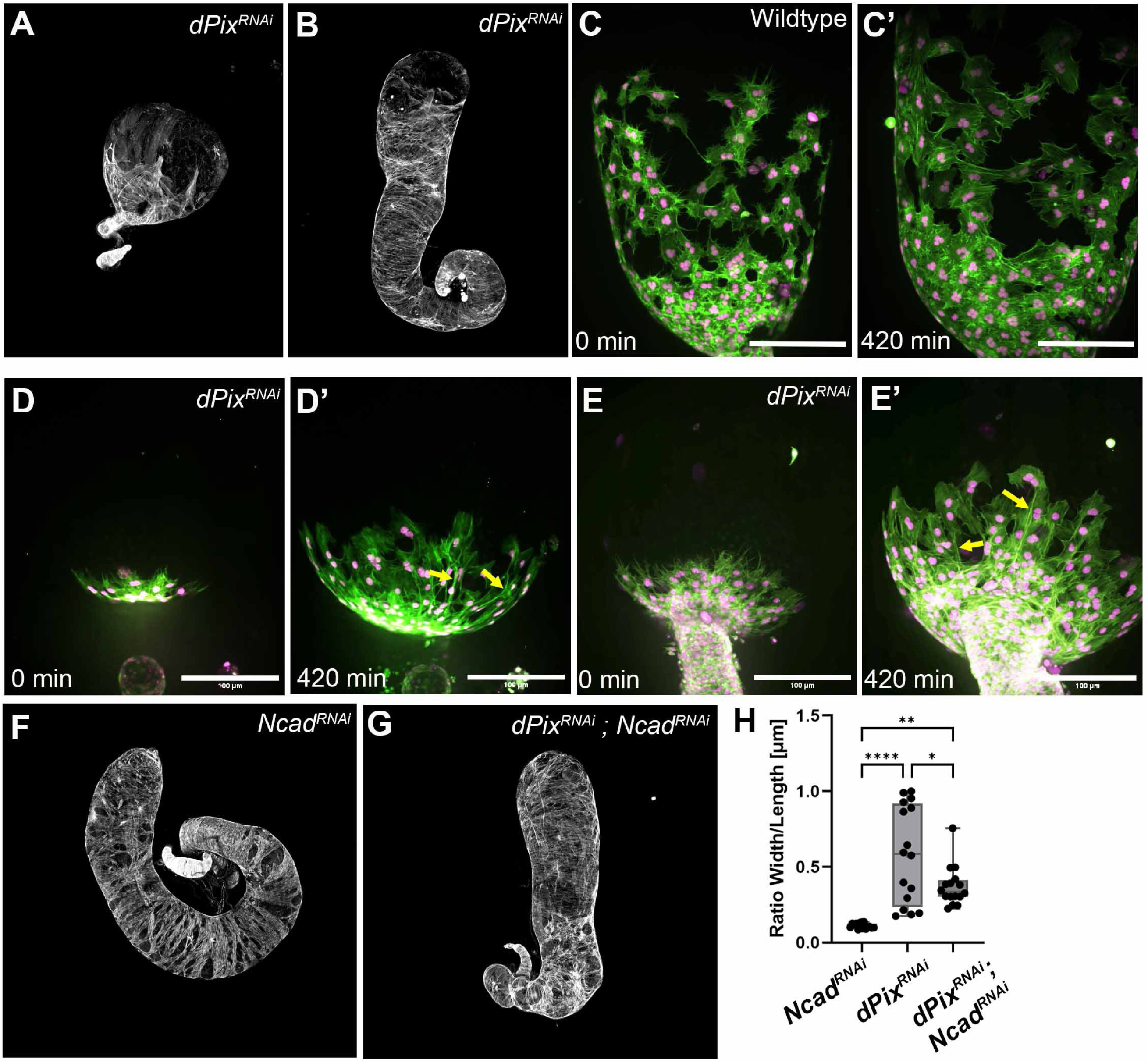
dPix knockdown dramatically slows myotube migration, and reducing N-cadherin partially suppresses the testis defects. A,B. Adult testis illustrating the range of phenotypes seen after *dPix^RNAi^*. C. Wildtype myotube migration. D, E. Rpresentative examples of the delay in migration after *dPix^RNAi^*. Some myotubes that are far apart remain interconnected by long processes that stretch over myotubes located between (arrows). F. Adult testis after *Ncad^RNAi^*. G. Adult testis after *dPix^RNAi^*; *Ncad^RNAi^*. H. Quantification of phenotypic severity.

To understand how migration was affected after *dPix^RNAi^*to lead to this severe adult phenotype, we live-imaged explanted pupal testis during migration. In wildtype, when nascent myotubes from the genital disc encounter the germline, they begin to migrate into the space between the underlying germline cells and the overlying pigment cells (Fig. 7C). Cells migrate as a mesenchymal cohort, joined by N-cadherin junctions. Our previous work revealed that cells are stimulated to migrate in the direction where they sense a free edge, with this directionality determined in part by slower turnover of focal adhesions at free edges and faster turnover at cell-cell borders (Bischoff *et al*., 2021). However, cells remain in contact with neighbors throughout migration (Fig. 7C, C’), and gaps between cells are closed by a combination of N-cadherin mediated adhesion and actin-based purse strings that form around gaps.

When we live-imaged *dPix^RNAi^* testis, we observed a dramatic change in migration behavior. Nascent myotubes moved onto the testis from the genital disc, but their progress from the proximal end was exceptionally slow (Fig. 7D,E; representative of eight movies). This is consistent with the adult phenotype, where only the most proximal tstis is covered in muscle. In fact, the movies we took understate the difference. We could not capture images from wildtype testis in which the myotubes were just starting to enter the testis, as at that stage in wildtype the connection between the genital disc and the testis was not fully formed, and they detach upon dissection—this suggests the testes we observed after dPix knockdown were in fact developmentally further along than one would think given the distance the myotubes had migrated. However, despite the delayed migration the cells did not dramatically differ from wildtype myotubes—they were well spread on the underlying cyst cells and had leading edge filopodia. One feature was notable; some myotubes that are far apart remain interconnected by long processes that stretch over myotubes located between (Fig. 7D,E, arrows), suggesting defects in cell-cell contact disassembly upon neighbor exchange. Thus *dPix^RNAi^* dramatically alters migration.

One possible explanation for the *dPix^RNAi^* phenotype was an effect on cell-cell adhesion. Perhaps elevated cell adhesion was preventing cells from escaping the proximal region. If this was the case, we reasoned that reducing cell-cell adhesion might suppress the *dPix^RNAi^* phenotype. Myotubes are connected by N-cadherin-mediated cell junctions, and adhesion is important for sealing gaps in the muscle sheet. Knockdown of N-cadherin leads to gaps in muscle coverage all along the proximal to distal axis (Fig. 7F; (Rothenbusch-Fender *et al*., 2017)). We thus asked if reducing N-cadherin by *Ncad^RNAi^* suppressed the *dPix^RNAi^*phenotype. As noted above, the phenotype of *dPix^RNAi^* varies from very strong reduction in muscle coverage and an almost round testis to examples where the testis was shortened in the proximal-distal axis but muscle coverage defects were less severe. To quantify suppression, we assessed the degree of shortening of the testis, calculating the ratio of width to length. *Ncad^RNAi^* alone does not significantly shorten the testis (Fig. 7H; n=16 for all treatments). *dPix^RNAi^* leads to a broad range of phenotypes, with some testis severely shortened and rounded and others less shortened (Fig. 7H). In contrast, animals in which both were knocked down (*dPix^RNAi^ Ncad^RNAi^*) had testis that were almost all in the less-severe category (Fig. 7G, H). Thus reducing cell-cell adhesion alleviates the effect of *dPix^RNAi^* .

### Knocking down different dPIX isoforms has different effects on testis shaping

The *dPix* gene is complex, with 8 differentially spliced isoforms (Fig. 8A). The catalytic Rho-GEF (DH) domain, the N-terminal SH3 domain and the PH domain are all encoded in the common exons shared by all isoforms (Fig. 8A). However, other exons differ in complex ways between the different isoforms. One major difference is between isoforms F, H, and I and the other isoforms. F, H, and I share a long exon just downstream of the catalytic domain that is missing in the other isoforms (Fig. 8A). This difference also means splicing of this FHI-specific exon into the next downstream exon is in a different protein reading frame, and thus all three of these also lack the downstream protein region containing the binding site for the Pix partner Git (Fig. 8A). Below we refer to these collectively as the FHI isoforms.

**Fig 8.**
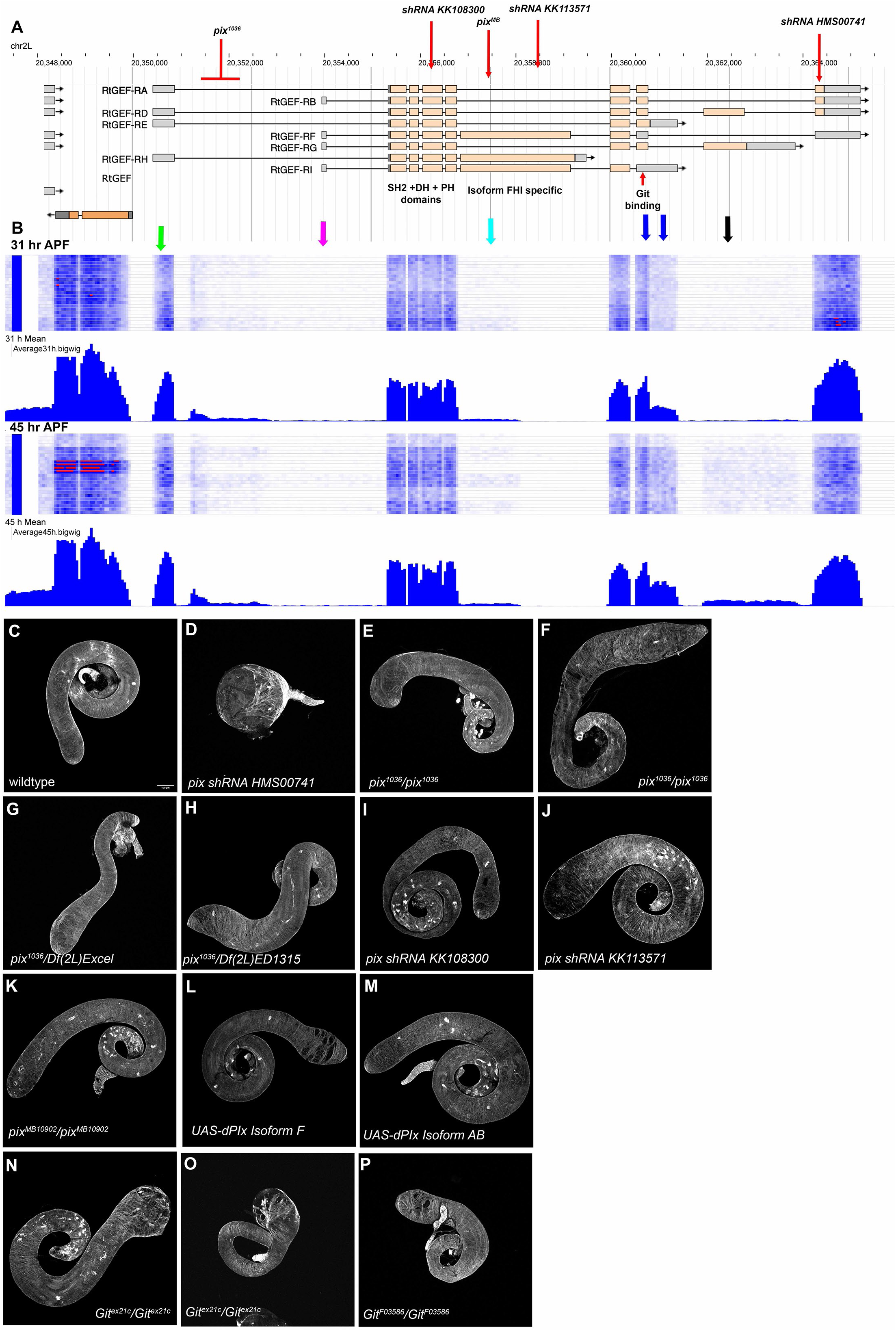
Different dPix isoforms play different roles in testis morphogenesis and the dPix binding partner Git also plays a role. A. Diagram of the genomic structure of the *dPix* gene, scale at top, 5’ end left, exons are grey (non-coding) or tan (protein coding) boxes and introns lines. Multiple *dPix* isoforms are illustrated. Above at the locations of three shRNAs targeting different exons and the location of mobile element insertions in two mutant alleles. Below are some features of the dPix protein isoforms. B. RNAseq data from the 31 hrAPF and 45 hr APF timepoints. Colored arrows indicate exons discussed in the text. C-O. Adult testes from wildtype, adults expressing the noted UAS-driven RNAi line targeting *dPix* under control of *mef-*GAL4, or the *dPix* or *Git* mutant alleles indicated. All are stained with fluorescently-labeled phalloidin to reveal F-actin, which highlights muscles. Phenotypes are discussed in the text.

Our RNAseq dataset allowed us to define which isoforms are likely to be expressed during myotube collective migration. When we mapped sequence reads to the map of different isoforms, we saw something surprising. At the 31 hr mid-migration time point, almost all transcription appears to start at the upstream start site (Fig. 8B, green versus magenta arrows), suggesting isoforms B, F, and G are expressed at very low to zero levels. There are, at most, very low levels of expression of the exon shared by isoforms D and G (Fig. 8B, black arrow), although these isoforms are detected in the 45 hr postmigration sample. This leaves isoforms A, E, H, and I. The long added exon shared by the FHI isoforms is also expressed at very low levels (Fig. 8B, cyan arrow),and intriguing the reads only extend part of the way through the predicted exon. Putting these data together, it appears that the predominant dPix isoforms expressed during migration are isoforms A and E, with levels of A about twice as high (Fig. 8B, blue arrows). Both of these share the predicted GIT—binding site (Fig. 8A).

Previous work revealed that different isoforms of dPix differ functionally, having differential importance in distinct tissues. The RNAi line which we used, *shRNA-HMS00741*, is the one used by Dent et al. (Dent *et al*., 2019) and it targets isoforms A, B, D, and F. They found that this RNAi line reduces levels of *dPix* mRNA and alters Hippo signaling in imaginal discs in ways similar to the effects of the strong allele *dPix^1036^*, suggesting that isoforms A, B, D, and F are important in those tissues. Isoform A, one of those targeted by this RNAi line, is also sufficient to rescue *dPix* mutant defects in oogenesis (Dent *et al*., 2019). Thus, the RNAi line we used would target isoform(s) important for function in these tissues.

However, while examining the role of dPix in neuromuscular synapse growth Ho and Treisman revealed additional complexity (Ho and Treisman, 2020),. In this tissue, knocking down isoforms A, B, D, and F, using the RNAi line we used, did not mimic the effects in that tissue seen the strong allele *dPix^1036^*. Instead, in that tissue, an RNAi line targeting the large exon shared by isoforms FHI (*shRNA-KK13571)*, and an allele with a mobile element insertion into this exon *dPix^MB10902^*, mimicked the effect of the strong allele *dPix^1036^*, suggesting that in this tissue the FHI isoforms are predominant. Most intriguing, when they used the RNAi line we used, *shRNA-HMS00741,* targeting isoforms A, B, and D, it had an effect on synapses opposite that caused by knockdown of isoforms F, H, and I (Ho and Treisman, 2020). From this and other data they concluded that FHI and ABD isoforms have an antagonistic relationship with one another.

To begin to explore the roles of different isoforms of dPix in shaping the testis, we used additional genetic reagents. As we outlined above, the RNAi line we identified in the screen, which knocks down isoforms A, B, and D, causes a very strong defect in migration, with most of the testis uncovered by muscle (Fig. 8C vs. D). We next examined animals homozygous for the strong allele *dPix^1036^* , which results from a mobile element insertion in the first intron (Fig. 8A) which has strongly reduced levels of protein in mutant embryos (Parnas *et al*., 2001). A subset of these animals the testis had no muscle coverage, suggesting failure of attachment of the genital disc to the testis (7/17 testis). In those with muscle coverage, the testis had dilated tips of variable severity (Fig. 8E,F; 10/17 testes). Animals with the strong allele *dPix^1036^*over a Deficiency that removes the *dPix* gene (Df(2L)Exel6046) had stronger but variable testis shape phenotypes, varying from a dilated tip to more extreme defects in testis shape (Fig. 8G; representative of 14 testes)—all also had reduced testis coiling. We observed similar defects in testis shaping in animals *dPix^1036^/Df(2L)ED1315* (Fig. 8H; representative of 13 testes). Finally, an RNAi like that should target all isoforms (*shRNA-KK108300*) also had dilated tips (Fig. 8I; representative of 4 of 7 testes). These data support a role for dPix in shaping the testis. However, intriguingly, knocking down isoforms F, H, and I, using the same RNAi line that gave Ho and Treisman their strongest phenotype (*shRNA-KK13571)*, had only mild effects on testis shape (Fig. 8J; 3 of 7 testes had dilated tips). *pix^MB10902^,* the insertional mutant that disrupts the shared exon of these isoforms, had no effect on testis shape (Fig. 8K; 5/8 testis examined), though a subset of mutants had apparent failure of attachment of the genital disc to the testis (3/8 testis examined). We thus tested the idea that FHI isoforms act antagonistically to the ABD isoforms. To do so, we used UAS-driven constructs that express the F-isoform. This led to penetrant distal tip expansion and gaps in distal muscle coverage (Fig. 8L; 18/22 testes examined). In contrast, overexpressing the AB isoform had little or no effect (Fig. 8M; 7 testes examined). Together, these data support the idea that the ABD isoforms play the largest role in the testis, consistent with the fact that isoform A is most highly expressed at the RNA level. They also are consistent with the idea of antagonism between different isoforms. To further explore this, we examined whether combining a mutant affecting the FHI isoforms, *pix^MB10902^,* with our *pix RNAi* targeting isoforms ABDF, might suppress the RNAi phenotype. However, while *pix^MB10902^* is homozygous viable and expressing that RNAi line alone does not lead to lethality, *pix^MB10902^* is lethal in combination with our strong *dPix-RNAi*, suggesting that balance among the different dPix isoforms is important for additional processes in other tissues.

### The dPix binding partner GIT plays an important role in testis shaping

PIX proteins are unique among RhoGEFs in that they can form heterodimers with GIT proteins, a family of Arf GAPs. Each can homodimerize or they can heterodimerize to form the PIX-GIT signaling scaffold. Some Pix roles are shared with GIT and others are not. For example. in Drosophila Pix and GIT work together in Hippo signaling in imaginal discs (Dent *et al*., 2015) and in regulating follicle cell epithelial architecture during oogenesis (Dent *et al*., 2019). In contrast, Git does not play a role in neuromuscular synapse growth (Ho and Treisman, 2020). Strikingly, the tissue where Git does not parallel Pix in function is one in which isoforms FHI, which lack the Git interacting region, are important, whereas tissues that evidence suggests rely on other isoforms like isoform A do require Git.

Since isoform A and most of the other isoforms containing the GIT binding site were targeted by the *dPix* RNAi line that gave our strong phenotype, and since those isoforms were also those most strongly expressed in migrating myotubes, we examined the effect of *GIT* mutants on testis shaping. We examined two different mutants, *git^Ex21c^*, a probable null allele that deletes the first 109 amino acids (Bahri *et al*., 2009), and *git^F03586^*, a piggyBac transposon insertion in an intron that is reported to be protein null (Podufall *et al*., 2014). *git^Ex21c^*homozygotes had penetrant defects, with strong enlargement of the distal testis and muscle coverage gaps (Fig. 8N,O; 14/17 testis examined). *git^F03586^* homozygotes also had distal enlargement and mscle coverage or alignment issues, though these were less penetrant (Fig. 8P; 4/8 testis examined). Thus, the Pix/Git complex regulates testis shaping.

### *dPix* RNAi elevates rather than decreases the activity of a cdc-42 sensor

Biochemical and cell biological studies revealed that mammalian beta-Pix can act a guanine nucleotide exchange factor (GEF) for Rac1 and Cdc42. We thus hypothesized that knockdown of dPix would reduce levels of cdc42 activity. We used an EGFP-tagged biosensor for cdc42 activity, based on the cdc42-interacting CRIB domain of the Pak family kinase Mbt. In previous work it was verified as functional by pulldown assays (Rötte *et al*., 2024). We extended this by testing it in testis myotubes expressing it there using the *mef-GAL4* driver. We analyzed myotubes on the seminal vesicle at the base of the testis where cortical localization was most easily visualized. The senor was strongly enriched at the cell cortex (Fig. 9A), consistent with the idea that endogenous cdc42 and/or Rac activity are high there. *cdc42* RNAi substantially reduced the cortical signal (Fig. 9B), something we verified using line scans across the membrane (Fig. 9C) and which we observed with two different *cdc42* RNAi lines (Fig. 9F). Rac2 RNAi also reduced the cortical membrane signal (Fig. 9D, E), though not by as much as by *cdc42* RNAi (Fig 9F).

**Fig. 9.**
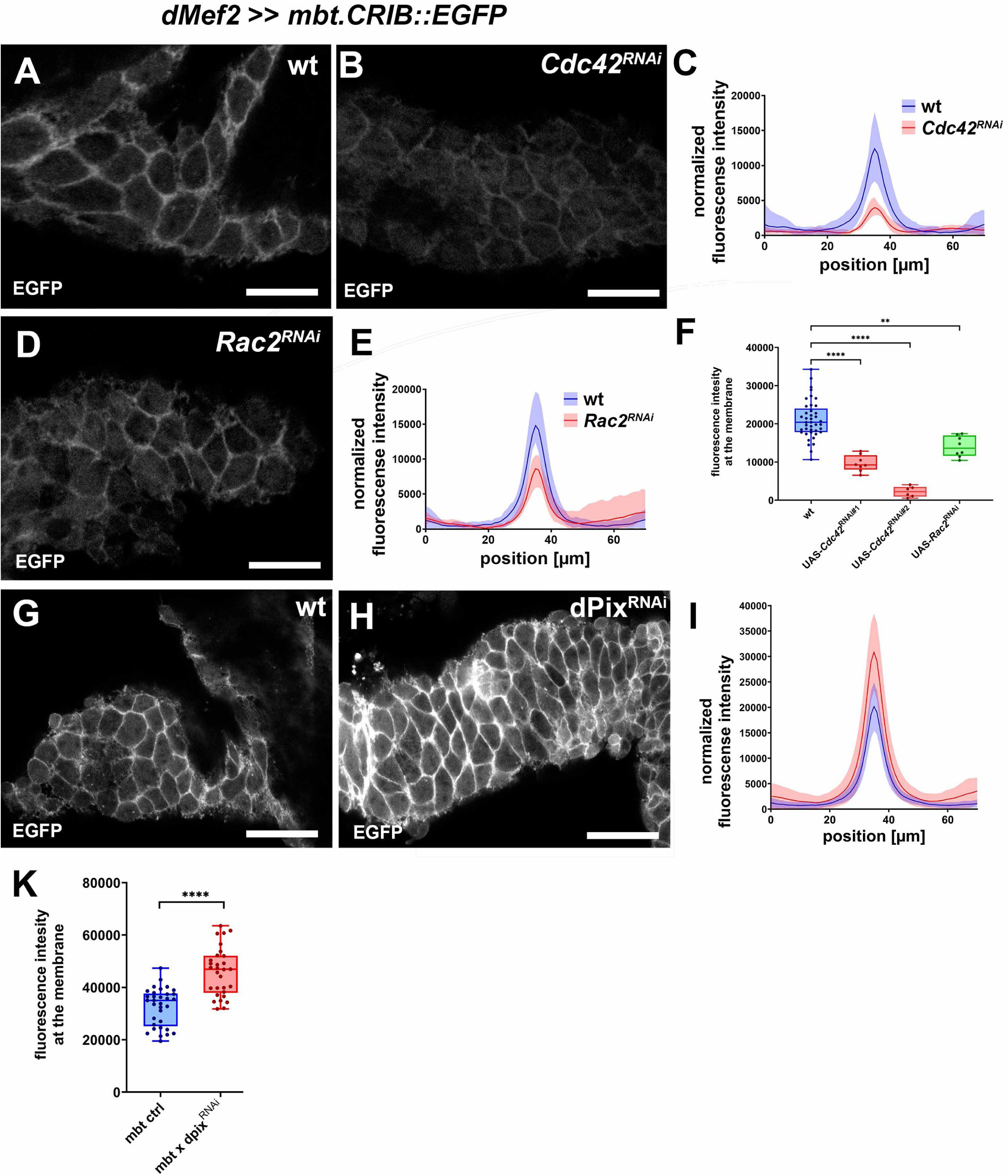
Knockdown of the AB isoforms of dPix elevates rather than reduces the signal froma cdc42 biosensor. A,B,D,G,H. Images of the cdc42 biosensor (the CRIB domain of Mbt fused to EGFP) expressed in the myotubes on the seminal vesicle at the base of the testis, using *dMef2*-GAL4. A. The biosensor signal is found at the cell cortex. B. *cdc42* knockdown reduces cortical biosensor signal. C. Line scans perpendicular to the membrane reveal the reduction in the cortical signal. wt: N=7, RNAi: N=8. D, E. Slightly reduced biosensor signal after Rac2 knockdown. wt: N=6, RNAi: N=8. F. Quantification of normalized membrane signal in wildtype, two different *cdc42* RNAi lines and a *Rac2* RNAi line. G,H. dPix knockdown elevates the cortical biosensor signal. wt: N=17, RNAi: N=30. I. Line scans perpendicular to the membrane. J. Quantification of normalized membrane signal.

We next knocked down dPix, using the RNAi like that has the strongest effect on testis migration. To our surprise, biosensor signal was not reduced as we had expected. Instead, it was increased somewhat (Fig. 9G vs H; representative of 17 wildtype and 30 *dPix^RNAi^* samples). Once again, this apparent elevation of the sensor at the cortex was verified by line scans across the membrane (Fig. 9I), and the difference in membrane signal was statistically significant across our samples (Fig. 9J). Together with the analysis above, this suggests different dPix isoforms may have surprising effects.

## Discussion

One key issue for our field is defining the molecular mechanisms underlying organogenesis. The Drosophila testis is shaped by a set of circumferential muscles that arise via the collective cell migration of myotubes from the genital disc onto the testis during pupal development and subsequently migrate to fully enclose the underlying germline and soma (Rothenbusch-Fender *et al*., 2017; Bischoff and Bogdan, 2021). We previously characterized this migration, revealing that these cells migrate in the confined space between the germline and the overlying pigment cells (Bischoff *et al*., 2021). These cells are loosely connected by N-cadherin mediated adhesion and move toward free edges, thus driving cells forward. They also close gaps between the cells, preventing later holes in muscle coverage.

### Intriguing parallels between collective cell migration and axon guidance

Our initial goal in setting up this screen was to identify regulators of collective cell migration and cell-cell adhesion, by identifying adult testis phenotypes we predicted would be caused by defects affecting these mechanisms. Many of our RNAi knockdowns fit the expected profile. These included several with partial coverage of the adult testis in muscle, leading to gaps in the distal region, consistent with defective migration. Others had gaps in muscle coverage all along the proximal-distal axis, suggestive of defects in adhesion. Our RNAseq data and the results of our knockdown screen provide some potentially interesting insights.

As neurons send out axons, they interact with other axons, with other cell types, and with extracellular substrates along their path in a form of directed cell migration. We found that many proteins known for their roles as axon guidance factors in the nervous system are highly expressed during testis nascent myotube collective cell migration. Further, knockdown of four of these cause defects in testis morphogenesis: *Plexin A*, *Netrin B, beat-IIIc*, and the Latrophilin homologue *Cirl*. In principle, axon guidance shares many features with contact-regulated modes of mesenchymal collective cell migration, with related emergent behaviors regulated via direct contact or secretion of guidance factors. Contact-dependent axon repulsion, for example, has features in common with contact inhibition of locomotion (Stramer and Mayor, 2017). During *Drosophila* follicle cell rotation, for example, Semaphorin/PlexA and Fat2/Lar-signaling are crucial to regulate contact-dependent planar-polarized protrusion formation, thus driving and synchronizing cellular locomotion (Stedden *et al*., 2019). Substrate-derived guidance cues are another feature that was previously recognized to be highly similar between axon guidance and collective cell migration (Aberle, 2019). Consistently, Netrin-signalling also affects collective cell migration, *eg.* in cultured mammalian liver cells (Han *et al*., 2019) and in vivo in *Drosophila* cardiac cell migration (Raza and Jacobs, 2016). One speculative possibility is that an ancestral function of these proteins is to regulate collective cell migration outside of the nervous system. Consistent with this notion, both Plexins and Netrins appeared early in or even before animal evolution and predate the evolution of the nervous system (Junqueira Alves *et al*., 2021; Cortes *et al*., 2023). The results of our screen provide exciting opportunities for future studies to define non-neuronal roles of canonical “axon guidance factors” by understanding their functions in the context of testis myotube migration.

### The regulation of Rho family and Ras/Rap GTPases plays important roles in many aspects of testis morphogenesis

The small Rho-family GTPases Cdc42, Rac2 and RhoA all play non-conventional roles in myotube collective cell migration. Cdc42 and Rac2 regulate integrin-adhesion lifetime and hence the ability to migrate, and RhoA stimulates retraction of cell edges and filopodia (Bischoff *et al*., 2021). The receptor tyrosine kinase Heartless (Htl) also plays a role, suggesting that Ras family small GTPases may be involved (Rothenbusch-Fender *et al*., 2017). One of our goals was to begin to uncover the regulators of these GTPases. To do so, we analyzed the consequences of knockdown of all *Drosophila* Rho-family and Ras/Rap GEFs and GAPs. This revealed that multiple GEFs and GAPs are necessary for normal testis morphogenesis and full muscle coverage. This suggests a high level of fine-tuning of GTPase activation and deactivation.

To illustrate this, we began to dissect the role of dPix in more detail. dPix is the fly homolog of mammalian beta-Pix, which is a guanine nucleotide exchange factor (GEF) for Rac1 and Cdc42. beta-Pix regulates diverse cellular processes from synaptogenesis to collective cell migration, both in vitro (Plutoni *et al*., 2016) and in vivo (Omelchenko *et al*., 2020). One of the strongest hits in our knockdown screen came when using a well-validated RNAi reagent (Dent *et al*., 2015) targeting dPix. This led to strong to nearly complete loss of muscle coverage of the adult testis. When we examined pupal testis myotube migration live, we found very strong delays in migration, consistent with this adult defect. However, as we dug deeper, the story became more complex. Pix has multiple isoforms and its function in different tissues appears to differentially depend on their respective functions. Our initial RNAi line targets a subset of the isoforms, A,B,D, and F. Dent et al. used this same RNAi line and found an important role for these isoforms in regulating Hippo signaling in imaginal discs (Dent *et al*., 2015). However, Ho and Treisman, studying growth of neuromuscular synapses, found something quite different. In this tissue the F, H and I isoforms play the key role, while all other isoforms act as antagonists of these isoforms (Ho and Treisman, 2020). Our findings suggest that both sets of isoforms have functions in testis shaping, and are consistent with the idea of antagonism, as overexpression of the F isoform led to defects in muscle coverage. We also found a role for the Pix binding partner Git in testis shaping—this made sense as Git can bind isoform A, which is the major isform expressed in the tests, and cannot bind isoforms FHI Future studies will have to define how the dPix-Git interaction relates to the differential Isoform-functions, and whether this somehow mediates dPix function as a GEF. Finally, we did two experiments to begin to assess the mechanism by which dPix knockdown alters migration. One hypothesis was that knockdown leads to too much cell-adhesion. We tested this by knocking down dPix and N-cadherin, hypothesizing that if we reduced adhesion, we might ameliorate the *dPix^RNAi^*phenotype. N-cadherin knockdown alleviated the effect of *dPix^RNAi^*, consistent with the idea that *dPix^RNAi^* may cause elevated cell-cell adhesion. In parallel, we tested the hypothesis that dPix knockdown would reduce the activity of a validated cdc42 sensor. To our surprise, it did not—in fact activity was elevated. This suggests that the dPix isoform was targeted is not simply acting as a cdc42 GEF. It will be interesting to explore how the different isoforms interact to shape the activity of small GTPases.

### Sculpting the testis is a multistep process and our screen provides leads into many different aspects of this

While we initially sought to identify regulators of migration and cell adhesion, the results of our screen opened our eyes to other aspects of testis organ morphogenesis. We found numerous candidates whose knockdown allowed full muscle coverage but still caused severe defects in testis morphology. We can now put these into the context of the shape changes of the testis we characterized after testis nascent myotubes migration (Bischoff and Bogdan, 2023). In that work, we identified distinct phases of morphogenesis (Fig. 10). The first phase is migration (30-36 hr APF) during which myotubes migrate from the genital disc to cover the testis. The next phase is myotube elongation, in which the myotubes elongate perpendicular to the proximal distal axis, beginning to take on the morphology of muscles (45 hr APF). The third phase is myotube condensation, in which the elongated myotubes narrow drastically and are linked by filopodial processes (53 hr APF). The final phase is proximodistal spreading and bilateral constriction, in which the muscles take on their mature circumferential arrangement and elongate the testis (66 hrs APF).

**Fig 10:**
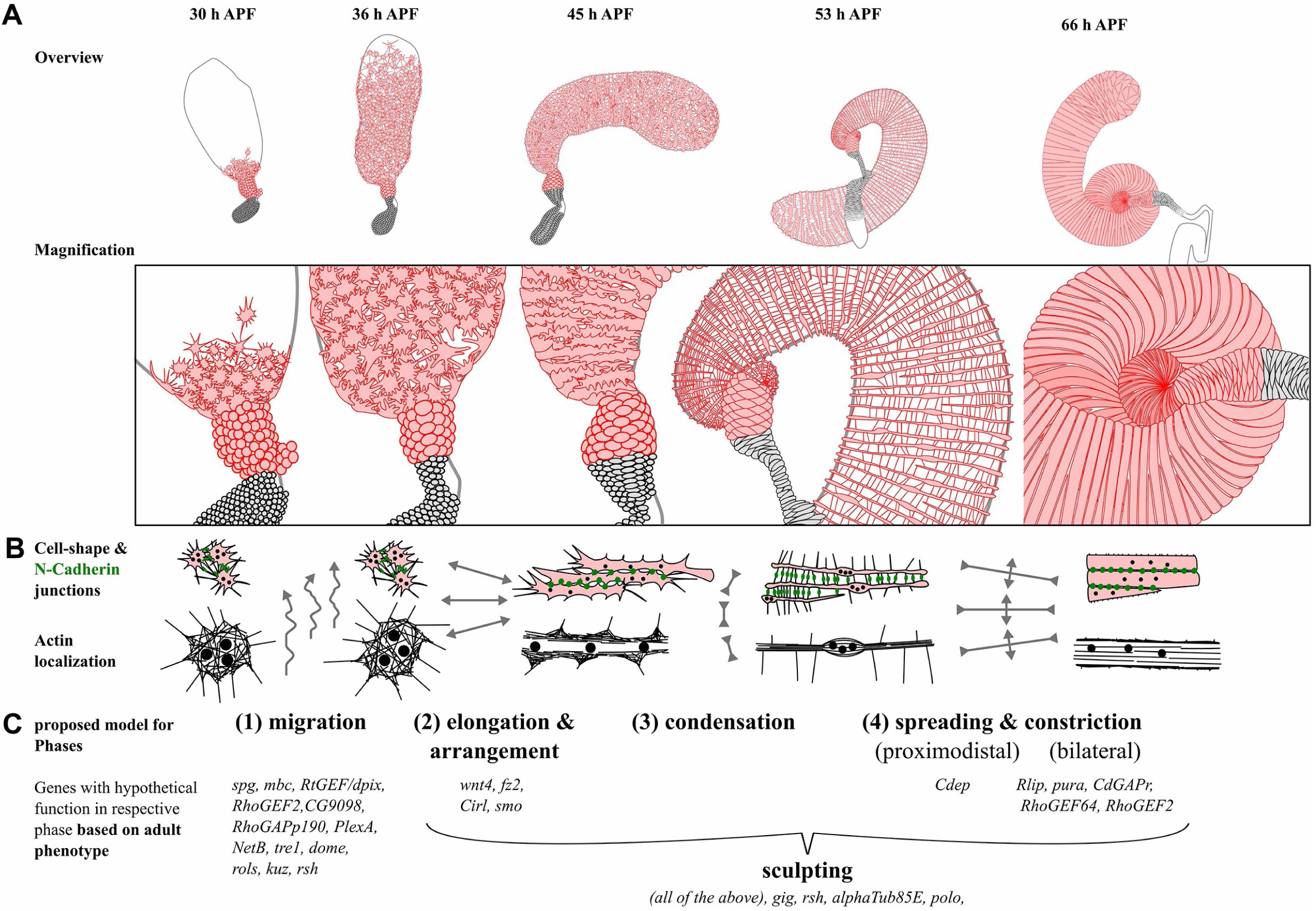
Cartoon illustrating the sequential events of testis morphogenesis and the steps at which we hypothesize each gene may act. A. Diagrams of the entire testis and closeups of the myotubes at each step. B. Changes in actin and N-cadherin localization at each step. C. Stages of morphogenesis and proposed steps at which different gens act.

Based on that analysis, we are now able to hypothesize which specific step or steps of development are likely affected upon knockdown of our positive screen candidates (Fig. 10C). Candidates that cause partial coverage or gaps in the sheet like *PlexA* and *mbt* likely affect the first migration phase, as this step was affected in earlier mutants we examined with coverage defects (Rothenbusch-Fender *et al*., 2017; Bischoff *et al*., 2021). The next crucial step in *Drosophila* testis-morphogenesis is the formation of the distinct and fascinating spiral-shape. Our findings suggest that this process relies on the correct myotube arrangement and alignment after migration. This seems to be self-regulated, as muscle-specific knockdowns that caused irregularities in the parallel organization of muscles also caused a loss of the spiral-shape. Furthermore, in Fz2 and Wnt4, we found a promising potential receptor/ligand pair that may be important for this self-regulation process. The profound defects in muscle arrangement caused by their knockdown suggest they might regulate the nematic ordering of myotubes.

Subsequently, as myotubes differentiate into muscle, they undergo an intriguing condensation and decondensation process, thinning and then re-expanding the myofibrils (Fig. 10B). Knockdown of *Cdep*, caused hyper-condensed muscle cells, suggesting a role in the de-condensation-process from 53 to 66 hrs APF. During the final de-condensation process, muscles become wider in the proximodistal axis (spreading), while constricting perpendicular to this axis, causing the testis to become thinner and even longer, thus increasing the number of revolutions in the spiral. Knockdown of multiple candidates, including *RhoGef2, pura, and RhoGEF64,* caused the entire testis – or parts of it – to become thinner or wider, suggesting a role of these GTPase regulators in this final phase of morphogenesis (Fig. 10C). Some candidates – like *gig* or *rsh* –causes defects, that cannot be categorized easily, but that cause shape-irregularities without affecting coverage, suggesting an important role during the sculpting process.

Together, the suite of different defects that emerged from our screen revealed that after migration the mechanical properties of the organ still must be fine-tuned, in an interplay between the muscles and the underlying tissue, to allow normal testis sculpting-morphogenesis. This is reminiscent of the shaping of the *Drosophila* oocyte by the follicle epithelium, which modifies its ECM to generate a molecular corset, enabling egg cell-elongation (Cetera and Horne-Badovinac, 2015). The process is also reminiscent of the fascinating sculpting of vertebrate airway epithelia by smooth muscle cells (Goodwin *et al*., 2019; Goodwin *et al*., 2023), and the many other cases in which smooth muscle cells sculpt other tissues during vertebrate development (Jaslove and Nelson, 2018) In the testis, the different contributions of the mechanical properties of the underlying ECM, the mechanical properties of the muscle cells themselves, and the role of the correct muscle cell arrangement remain to be elucidated, revealing their individual roles in correct sculpting. We’re excited to use this system to explore the underlying mechanisms, revealing how simple self-regulated organization can lead to tissue-sculpting via a process with intriguing morphological complexity.

## Acknowledgements

We are very grateful to the Bloomington and Vienna Drosophila Stock Centers for many shipments of RNAi lines, to Jessica Treisman and Sally Horne-Badovinac for Drosophila stocks, to Nat Prunet and the Biology Imaging Core for technical support, Roman Bandy of the UNC Flow Cytometry Core, Gabrielle Cannon of the UNC Advanced Analytics Core, the Peifer lab for helpful feedback on the manuscript, and the Peifer, Bergstralh/Finegan and Williams labs for feedback throughout. M.C.B was supported by the DFG Walter Benjamin Programme (ref. GZ: BI 2384/1-1) and work in the Peifer lab is supported by NIH R35 GM118096.

---

**Fig. S1.**
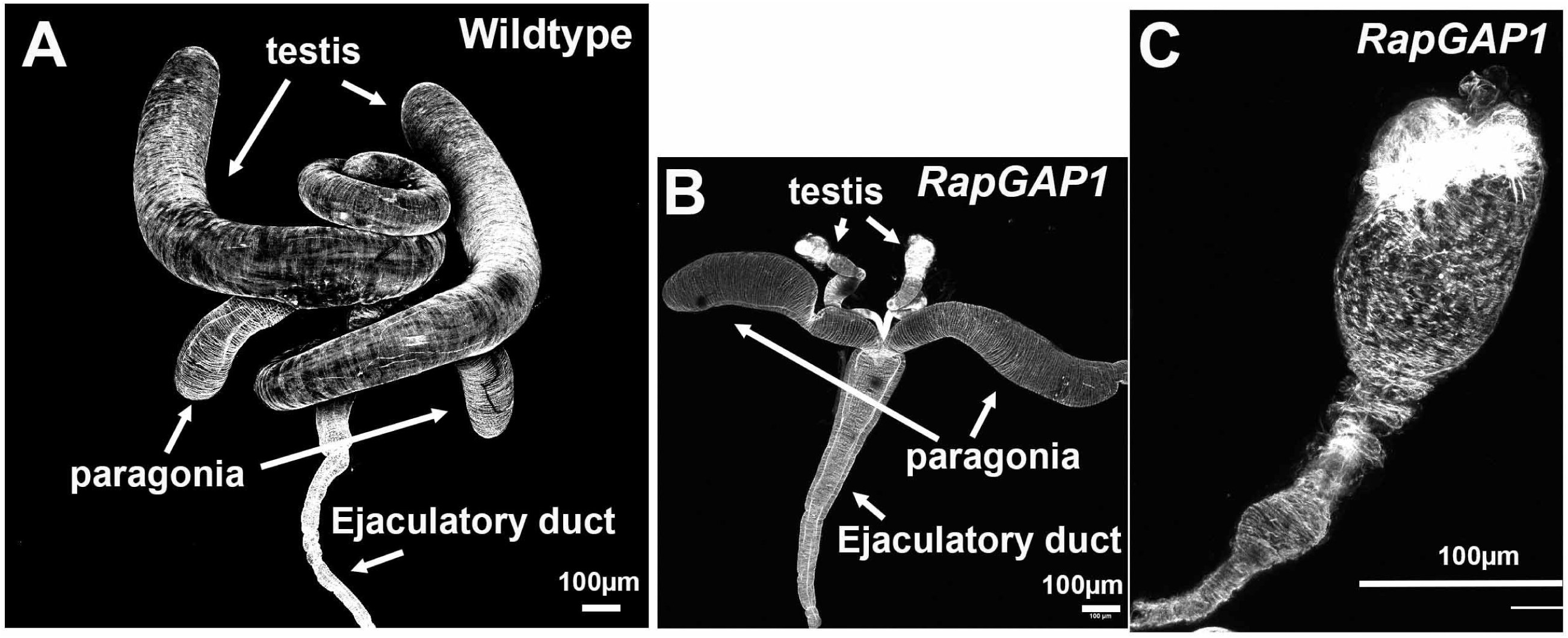
*RapGAP1^RNAi^*dramatically reduces the size of the testis. A,B. Overviews of the testis and associated somatic tissues in wildtype (A) and after *RapGAP1^RNAi^* (B). C. Closeup of the remnant testis after *RapGAP1^RNAi^*.

